# SEPSIS-INDUCED LIPID DROPLET ACCUMULATION ENHANCES ANTIBACTERIAL INNATE IMMUNITY

**DOI:** 10.1101/2025.01.29.635463

**Authors:** Filipe S. Pereira-Dutra, Julia Cunha Santos, Ellen Kiarely Souza, Rodrigo Vieira Savi, Tamyris S. Souza, Hugo Espinheira-Silva, Felipe Ferraro-Moreira, Guilherme Iack, Tamires Cunha-Fernandes, Tathiany Igreja-Silva, Lohanna Palhinha, Mariana Macedo Campos, Douglas Mathias Oliveira, Vinicius Soares Cardoso, Matheus A. Rajão, Livia Teixeira, Luciana Souza-Moreira, Maria Fernanda Souza Costa, Patrícia Alves Reis, Patrícia T. Bozza

## Abstract

Lipid droplets (LDs) are lipid-rich organelles recognized as central players in lipid homeostasis, signaling, and inflammation. While their functions in inflammation are well-documented, the role of LDs in antibacterial immunity and infection resistance remains less understood. In this study, we investigated triglyceride synthesis and LD accumulation in the context of antibacterial innate immunity during sepsis. Our results show that LD accumulation is part of immunometabolic reprogramming in *E. coli*-infected macrophages. Pharmacological inhibition or genetic knockdown of DGAT1, a key enzyme in triglyceride synthesis, reduced LD formation, bacterial clearance, and pro-inflammatory responses (nitric oxide, PGE_2_, CCL2, IL-6, IFN-β). Notably, DGAT1 inhibition impaired the expression of several interferon-stimulated genes (ISGs), including viperin, iNOS, cathelicidin, and IGTP, in *E. coli*-infected macrophages. In a sepsis model, DGAT1 inhibition reduced sepsis-induced LD accumulation in peritoneal cells and decreased levels of CCL2, IFN-β, nitric oxide, and lipid mediators (PGE_2_, LTB_4_, and RvD1). Furthermore, DGAT1 inhibition accelerated sepsis-related mortality, coinciding with elevated bacterial loads in the peritoneum and bloodstream at 6 and 24 hours post-sepsis. Our results demonstrate that tryglicerides synthesis and LDs are critical regulators of infection resistance, contributing to both bacterial clearance and the coordination of a protective proinflammatory response during sepsis.

## INTRODUCTION

Lipid droplets (LD) are dynamic lipid-rich organelles that play essential intracellular roles, including regulating cell lipid homeostasis, intracellular signaling, and inflammatory processes ^1^. LD accumulation is heavily influenced by the flux of cellular lipid metabolism, including both the uptake and synthesis of neutral lipids, as well as lipid-consuming processes like lipolysis and β-oxidation ^2^. The presence of LD in leukocytes is a structural marker of the inflammatory response ^3^, since this organelle is also an important platform for a broad-range synthesis of eicosanoids and cytokine signaling ^4^.

Numerous studies on pathogen-leukocyte interactions have demonstrated a strong link between lipid droplets (LDs) and bacterial infections ^5–7^. Activation of Toll-Like Receptors (TLRs) plays a central role in inducing LD accumulation in leukocytes ^8–11^. Additionally, bacterial virulence factors have been shown to drive LD formation during infections by pathogens such as *Salmonella enterica* Typhimurium, *Listeria monocytogenes*, and *Mycobacterium tuberculosis* ^12–14^. Intracellular bacteria may exploit LDs as an energy source and as a mechanism to evade the immune system ^5^. In pathogenic bacteria, LDs serve as platforms for prostaglandin E_2_ (PGE_2_) synthesis, which has been linked to increased IL-10 production and suppression of Th1 cytokines, promoting bacterial survival and/or proliferation ^15–17^.

The perception of LDs solely as pro-pathogenic factors in infectious and inflammatory contexts has recently been challenged ^18–20^. Emerging evidence suggests that LDs play a pivotal role in interferon (IFN) signaling pathways and the effectiveness of early innate responses ^21–23^. Recent data show that LDs is a central hub in innate immunity, responding to danger signals by reprogramming cellular metabolism and serving as autonomous platforms for innate immune responses ^24^. In this context, the presence of antibacterim peptide cathelicidin (CAMP) in LDs confers greater protection against bacterial infection in human macrophages ^24^. These findings position LDs as key regulators of metabolic reprogramming and facilitators of antimicrobial mechanisms ^24^. Furthermore, the involvement of LDs in the antibacterial response is evolutionarily conserved, being first reported in *Drosophila* ^25^.

Sepsis is a complex life-threatening syndrome caused by dysregulated inflammatory and metabolic host response to infection^26^. The balance between resistance to infection and tissue tolerance of it might be the key to understanding the molecular mechanisms of sepsis ^27,28^. Alterations in lipid metabolism and increased LDs are observed during sepsis ^11,29^, however although LD functions in inflammation are well-documented, the role of LDs in antibacterial immunity and infection resistance and their roles in sepsis survival remains less understood. During sepsis, LDs are critical for amplifying the proinflammatory response but also contribute to the disruption of tissue tolerance mechanisms leading to increased organ dysfunction ^29^. Notably, the pathways driving the breakdown of tissue tolerance may play a key role in infection resistance ^27,28^. In this study, we investigated the formation of LDs and the triglycerides synthesis in antibacterial innate immunity during sepsis. Our results demonstrate that LDs are critical regulators of resistance to infection, contributing to both bacterial clearance and the coordination of a protective proinflammatory response during sepsis.

## 1. Materials and Methods

### Reagents

Recombinant Murine interferon γ (IFNγ) was obtained from PeproTech. From Sigma-Aldrich (St Louis, MO) were obtained Lipopolysaccharides from *Escherichia coli* (serotype O111:B4, cat.# L4931), A922500 (Cat# A1737), CI976 (Cat# C3743), oleic acid (Cat# O1008), Saponin from Quillaja bark (Cat# S7900), and Oil Red O (Cat# O0625) were purchased from Sigma-Aldrich/MERK. The Luria-Bertani broth (Cat# K25-1551), and Tryptic Soy Agar (Cat# K25-610052) was obtained from Kasvi (São José do Pinhais, PR, Brazil). RPMI-1640 (cat# 22400-089), penicillin-streptomycin (Cat# 15140148), gentamicin (Cat# 15750060), and L-glutamine (Cat# 25030081) were obtained from Gibco (Grand Island, NY, USA). DAPI (Cat# D1306) and Fluoromount-G Mounting Medium (Cat# 00-4958-02) were purchased from ThermoFisher Scientific (Waltham, MA, USA).

### Mice

C57BL/6J male mice were supplied by the Institute of Science and Technology in Biomodels from Oswaldo Cruz Foundation and used at 8–12 weeks of age. The Institutional Animal Welfare Committee approved all animal experiments in agreement with the Brazilian National guidelines supported by CONCEA (Conselho Nacional de Controle em Experimentação Animal) under license number L025/15 and L005/20 (CEUA/FIOCRUZ). Mice were maintained with standard rodent diet (AIN-93 M) and water available *ad libitum* with 12h light-dark cycle under controlled temperature (23 ± 1 °C).

### Macrophages cell culture

To obtain bone marrow-derived macrophages (BMDM), bone-marrow cells isolated from femur and tibia of mice were cultured for 7 days in RPMI-1640 medium supplemented with 18% (vol/vol) L929-derived M-CSF conditioned medium, 20% (vol/vol) heat-inactivated fetal bovine serum, 1% L-glutamine (vol/vol), and 1% penicillin-streptomycin (vol/vol) as previously described by Assunção et al., (2017). Differentiation was performed at 37 °C in a humidified 5% CO_2_ incubator. After seven days, adherent macrophages were harvested and seeded for assays. Differentiated macrophages were cultured in RPMI-1640 supplemented with 10% heat-inactivated fetal bovine serum (FBS, vol/vol), 1% L-glutamine (vol/vol), and 1% penicillin/streptomycin (vol/vol). BMDM cells were seed at a density of 0.1x10^6^ – 1x10^6^ cells/ml, and the cell culture were performed at 37°C in 5% CO_2_.

### Bacterial strains and growth conditions

*Escherichia coli* (ATCC 25922) used in this study were obtained from the Enterobacteria Collection (CENT) of the Oswaldo Cruz Foundation. The bacteria were cultured in Luria-Bertani broth (LB) for 16–18 h at 37°C to obtain stationary growth phase cultures. Before infection, the bacteria were washed three times with Phosphate-Buffered Saline (PBS), centrifuge (1,000 × g) for 10 min at 4°C, and resuspended in PBS. The bacteria were resuspended in PBS and diluted in RPMI with FBS to an appropriate multiplicity of infection (MOI) according to the optical density (OD) at 600 nm. *E. coli* (K-12 strain) Texas Red™ conjugate BioParticles (Cat# E2863, ThermoFisher Scientific) were also washed three times with PBS, centrifuge (1,000 × g) for 10 min at 4°C, and resuspended in PBS.

### *E. coli* infection of macrophages and treatments

An *in vitro* gentamicin protection assay was used to measure the phagocytosis and intracellular survival of *E. coli* based on the method described by Lissner et al. (1983), with modifications of Souza et al. (2021). The macrophages were seeded at 1 x 10^5^ and 5 x 10^5^ cells per well in 24- and 12-well places (flat-bottom, tissue-culture-treated plates; Costar), respectively, and were incubated for 12 h. Meanwhile, *E. coli* cells were cultured overnight at 37°C with agitation. Macrophages were infected with *E. coli* at a MOI of 100 for 1 h. Then, the culture medium was discarded, and the cells were washed with PBS with 100 µg/mL gentamicin three times. RPMI-1640 supplemented with 100 µg/mL gentamicin was added to each well to kill non-phagocyted bacteria, and incubation was continued for another 1 h. After incubation, media with gentamicin were removed, and fresh media was added for the remainder of the time. A similar protocol was used for *E. coli* bioparticules stimulation at MOI of 100. Alternatively, BMDMs were stimulated with LPS serotype 0111:B4 (500 ng/mL) plus murine IFNγ recombinant (10 ng/mL) for 24h. All experiments were done in triplicate.

To increase the LD accumulation, 40 μM of oleic acid was added to BMDM cell culture and incubated for 16 hr before *E. coli* infection. To impair the LD biogenesis, DGAT1 inhibitor (A922500) or ACAT1 inhibitor (CI976) were added to the cell culture and incubated for 30 min before *E. coli* infection and remained for all the infection time. The inhibitors were dissolved in dimethyl sulfoxide (DMSO, Sigma).

For Colony-forming unit (CFU) analysis, BMDM cells were washed three times with PBS and lysed with 10% Triton X-100 solution at the indicated time points. The CFU of bacteria were counted by plating the appropriate dilution in TSA plates.

### Lipid droplets staining and quantification

BMDM were fixed with 3.7% formaldehyde for 10 min, and LDs were stained with 0.3% Oil Red O’ (diluted in isopropanol 60%) as previously described ^32^. Preparations were analyzed with a 60× objective in FluoView FV10i Olympus confocal microscope (Tokyo, Japan). Images were acquired, colored, and merged using Olympus FV10-ASW and open-source ImageJ software (https://imagej.nih.gov/ij/). The Oil red O-stained LDs were measured using the open-source ImageJ software.

### Measurement of Lactate

Lactate levels in cell-free culture supernatant were measured using an enzymatic lactate kit (Labtest, cat.# 138-1/50) performed as per manufacturer’s instruction.

### Oxide nitric assay

The nitric oxide (NO) levels in culture supernatants and peritoneal lavage supernatants were measured by the colorimetric Griess assay. Briefly, 25μL of the sample was mixed with an equal part of Griess reagent (1% sulfanilamide / 0,1% N-(1-naphtyl) -ethtylenediamine dihydrochloride in 2.5% H_3_PO_4_) in a 96-well plate. The color development was assessed at 450 nm with a microplate reader (Spectramax, Molecular Devices, Inc., USA). As a standard curve, a solution of sodium nitrite (NaNO_2_) was used at an initial concentration of 200µM, diluted in fresh culture medium.

### Quantitative PCR (qPCR)

RNA was extracted with QIAamp Viral RNA (Qiagen®) from cells seeded (10^5^ cells/well) in 24-well plates. Quantitative RT-PCR was performed using a dye-based GoTaq 1-Step RT-qPCR System (Promega, Fitchburg, WI, USA) in a StepOne™ Real-Time PCR System (Thermo Fisher, Carlsbad, CA, USA). Amplifications were carried out in 15 µL reaction mixtures containing 2× reaction mix buffer, 1x of probe-based oligos from predesigned TaqMan Gene Expression Assays (Thermo Fisher, Carlsbad, CA, USA), and 5 µL of RNA template. The program for probe-based amplifications was 10 min at 95 ◦C followed by 50 cycles of 15 s at 95◦C and 1 min at 60 ◦C. The relative mRNA expression was calculated by the 2− ΔΔCt method. The *β-actin* (*actb*) expression was used as a reference gene. The probe-based oligos were all Predesigned Taqman Gene Expression Assays: *fasn* (ref: Mm00662322, g1, FAM), *dgat1* (ref: Mm00515643_m1, FAM), *acat1* (ref: Mm00507463_m1, FAM), *plin2* (ref: Mm00475794_m1, FAM), *plin3* (ref: Mm04208646_g1, FAM), *atgl/pnpla2* (ref: Mm00503040_m1, FAM), *cd36* (ref: Mm01135198_m1, FAM), *abca1* (Ref: Mm00442646_m1, FAM), *il-1β* (ref: Mm00434228_m1, FAM), *il-10* (ref: Mm00439614_m1, FAM), *cox-2/ptgs2* (ref: Mm00478374_m1, FAM), *5-lo/alox5* (ref: Mm01182747_m1, FAM), 15-LO (Cat# Mm00507789_m1) *inos/nox2* (ref: Mm00440502_m1, FAM), *arg1* (ref: Mm00475988_m1, FAM), *chi3l3/ym1* (ref: Mm00657889_m1, FAM), *mr1* (ref: Mm00468487_m1, FAM), and *β-actin/actb* (ref: Mm02619580_g1, FAM).

### Flow cytometry

Macrophages (5x10^5^) were incubated with the appropriate concentration of purified rabbit polyclonal antibody anti-GLUT1 (Cat# 21829-1-AP, Proteintech, USA), Alexa Fluor® 488 rat anti-mouse CD206 (MMR) Antibody (Cat# 141710, Biolegend, USA), APC mouse anti-mouse CD36 (Cat# 562744, BD Biosciences, USA), APC hamster anti-mouse CD80 (Cat#560016), and/or PE rat anti-mouse F4/80 Antibody (Cat#123109, Biolegend, USA) for 30 min at 4°C, after incubation with 10% of FBS in PBS/0.1% Sodium Azide to block non-specific binding. For GLUT1 staining, Alexa546 goat anti-rabbit IgG was added to all wells and incubated for 30 min. As negative control cells received only secondary antibodies. At least 10^4^ cells were acquired per sample in FACSCalibur flow cytometer and CellQuest™ software (Becton Dickinson, San Jose, CA, USA). All data were displayed on a log scale of increasing fluorescence intensity and presented as histograms. Analysis was performed by using FlowJo™ Software (Becton Dickinson, San Jose, CA, USA).

### Sepsis Induction and Treatment

Sepsis was induced by Cecal Ligation and Puncture (CLP), according to Reis et al. (2017), with modifications of Teixeira et al. (2023). Briefly, C57BL/6 mice were anesthetized with intraperitoneal injections of ketamine (100 mg/kg) and xylazine (10 mg/kg) and a 1 cm incision was made on the abdomen. The cecum was exposed and linked below the ileocecal junction. It was made 4 perforations (severe sepsis model) using a 22-gauge needle, a small amount of fecal material was expelled into the peritoneal cavity, and the cecum was gently relocated. The area was sutured with nylon 3-0 (Shalon) in two layers. Sham-operated animals (control) with identical laparotomy but without ligation and punctures.

After 6h and 24h post-surgery, Sham and CLP mice were orally treated with A922500 at a dose of 3 mg/kg or vehicle. All mice received 500 µL of sterile saline with meropenem (10 mg/kg; Merck) subcutaneously as fluid resuscitation and antibiotic therapy at 6h and 24h hours after surgery. Animals were monitored for 48h for survival and clinical score analysis. The clinical score was determined by the observation of following signs: piloerection, curved trunk, alterations in gait, seizures, limb paralysis, coma, respiratory rate, skin color alterations, heart rate, lacrimation, palpebral closure, decreased grip strength, limb, abdominal and body tone and body temperature alterations. The clinical evaluation was based on a multifactorial SHIRPA protocol, with the modifications of Reis et al. (2017).

Six or twenty-four hours after sepsis induction, the animals had their peritoneal cavity opened and washed with 3 ml of sterile saline. The peritoneal lavage was collected for total and differential cell count, CFU evaluation, LDs, cytokine, chemokines, lipid mediators, and NO quantification. For CFU evaluation, peritoneal lavage from each animal was diluted and plated on tryptic soy agar (TSA) plates. After 24 hours of incubation at 37°C, the number of bacterial colonies was enumerated manually.

### Leukocyte Count and Lipid Droplet Staining

Peritoneal lavage samples were diluted in Turk’s solution (2% acetic acid), and the total cell counts were performed with optical microscopy in the Neubauer Haemocytometry chamber. For differential cell count and LD Staining, the samples were cytocentrifuged in a microscope slide and fixed with 3.7% formaldehyde for 10 min. Cells were washed three times in PBS, slides were stained with 0.3% Oil Red O’ (diluted in isopropanol 60%) for 2 minutes at room temperature. Cells were washed once in 30% isopropanol and three times in distilled water. Slides were rinsed and counterstained with Mayer’s hematoxylin for 3 minutes. After incubation, the slides were washed three times in distilled water. Mounting solution and coverslips were added. LDs were enumerated by optical microscopy in 50 consecutive leukocytes.

### Measurements of Cytokines and Chemokines

CXCL1 / KC, CCL2 / MCP-1, IL-1β, IL-6, IL-10, IL-12p40, TNF-α, IFN-β and IFN-γ in cell-free culture supernatants and peritoneal lavage were measured using mouse Duoset ELISA kit (R&D Systems, USA) according to manufacturer’s instructions. Cathelin-related antimicrobial peptide (Cramp) in peritoneal lavage were mouse Sandwich ELISA kit (Cat# CSB-E15061m, CUSABIO, USA). The level of IFN-alpha in cell-free culture supernatants and peritoneal lavage were measured using mouse IFN-alpha All Subtype ELISA Kit (Catalog #MFNAS0, R&D Systems, USA)

### Measurements of Lipid mediators of inflammation

According to the manufacturer’s instructions, levels of Prostaglandin E2 (PGE_2_), Leukotriene B4 (LTB_4_) and Resolvin D1 (RvD1) in macrophages supernatant and peritoneal lavage were measured using EIA kits (Cayman Chemical, USA).

### Lipid droplets purification and Bacterial Plate Assay

BMDMs were seeded at a density of 25 × 10^7^ cells/ flask (175 cm^2^). In the next day, cells were stimulated with LPS serotype 0111: B4 (500 ng/mL) plus murine IFN-γ recombinant protein(10 ng/mL) for 24 h, at 37°C in 5% CO_2_. LDs were isolated from 4 culture flasks for each condition as previously described by Samsa *et al*, 2009 ^34^. Cells were gently scraped in media and centrifuged in a swing out bucket rotor at 200*g* for 15 min. The cell pellet was washed twice in PBS and then resuspended in 2.5 mL of lysis buffer (20 mM Tris, 1 mM EDTA, 1 mM EGTA, 100 mM KCl buffer (pH 7.4), followed by incubation on ice for 5 min. Cell lysis was completed using a nitrogen cavitator by applying nitrogen gas at 750 psi for 5 min. The cell lysate was collected in 15 mL conical tubes. The total cell lysate was centrifuged at 900*g* for 15 min. The resulting postnuclear supernatant was collected and applied to a sucrose gradient (1.08 M, 0.27 M, and 0.135 M) and subjected to ultracentrifugation (150,000 x g for 70 min). After ultracentrifugation, the first fraction was rich in isolated LDs. TEE-KCl buffer was added to the first fraction, and the LDs were isolated a second time by centrifugation at 14,000 × g for 15 min. As a control for proper cell fractionation, the activity of the cytoplasmic enzyme lactate dehydrogenase (LDH; Promega, G1780) was assayed.

Colony formation assays were conducted to assess the antibacterial activity of LDs as described by Bosch *et al* (2020). Exponentially growing *E. coli* cultures in LB broth at 37 °C were harvested at an OD600 of 1. The cells were then centrifuged, washed, and resuspended in 10 mM Tris-HCl buffer (pH 7.4) supplemented with 1% (v/v) LB medium. Aliquots of 100 μL of this bacterial suspension were incubated with 100 μL of intact LDs prepared using a sucrose gradient. Following incubation, the samples underwent 6–8 serial 1:10 dilutions in 10 mM Tris-HCl (pH 7.4) and were plated in triplicate on LB agar plates. CFU were quantified after 16 hours of incubation at 37 °C.

### DGAT1 Knocking down assays

BMDMs were plated (2,5 x 10^5^ cells/well) in 24-well culture plates and incubated for 12 h at 37 °C and 5% CO_2_. Cells were transfected with 100 pmol/μL siRNA targeting DGAT1 (Cat# s64951, ThermoFish Scientific) or scramble RNA (Cat# 4390844, ThermoFish Scientific) in Opti-MEM (Gibco, USA), using Lipofectamine RNAiMax (ThermoFish Scientific). Alternatively, macrophages were also transfected with 5µM self-deliverable AUM*silence* oligos for DGAT1 (Cat# AUM-SIL-A-100-Dgat1-1) or scramble RNA (AUM-SCR-A-100) according to the manufacturers’ instructions (AUM BioTech, LLC, USA). After 48 hours of recovery, BMDMs were then infected with *E. coli*. The efficiency of *knocking down assays* was measured 48-72 h post-transfection through quantitative RT-PCR performed using dye-based GoTaq 1-Step RT-qPCR System.

### Western blotting

BMDM cells were harvested using ice-cold lysis buffer pH 8.0 (1% Triton X-100, 2% SDS, 150 mM NaCl, 10 mM HEPES, 2 mM EDTA containing protease inhibitor cocktail—Roche). Cell lysates were heated at 100°C for 5 min in the presence of Laemmli buffer pH 6.8 (20% β-mercaptoethanol; 370 mM Tris base; 160 μM bromophenol blue; 6% glycerol; 16% SDS). Twenty μg of protein/sample were resolved by electrophoresis on SDS-containing 10% polyacrylamide gel (SDS-PAGE). After electrophoresis, the separated proteins were transferred to nitrocellulose membranes and incubated in blocking buffer (5% nonfat milk, 50 mM Tris-HCl, 150 mM NaCl, and 0.1% Tween 20). Membranes were probed overnight with 1:1000 dilution of the following antibodies: anti-Cathelicidin (#ab180760), anti-IRGM3 (#14979S), anti-Viperin (#13996S), anti-iNOS (#ab15323), anti-PLIN-2 (#15294-1-AP), and anti-β-actin (#66009-1-Ig). After the washing steps, they were incubated with IRDye—LICOR or HRP-conjugated secondary antibodies. All antibodies were diluted in blocking buffer. The detections were performed by Supersignal Chemiluminescence (GE Healthcare) or by fluorescence imaging using the Odyssey system.

### Statistical analysis

Data obtained in this study were presented as mean ± SEM of three to six independent experiments. The impaired two-tailed t-test was used to evaluate the significance of the two groups. Multiple comparisons among three or more groups were performed by one-way ANOVA followed by Tukey’s multiple comparison test, for all analyses was adopted p ≤ 0.05 as considered statistically significant.

## 2. RESULTS

### *Escherichia coli* induced-LD accumulation is part of pro-inflammatory reprogramming of murine macrophages

LD accumulation in leukocytes is a well-documented phenomenon both in experimental models of sepsis as well as in septic patients ^35,36^. However, the role of LD during infections caused by extracellular bacteria has not been fully clarified. Our first step was to investigate the participation of LD in the antibacterial response of macrophages. For it, we infected primary macrophages with *Escherichia coli*, a classical extracellular bacterium. The *E coli* infection led to a significant LD accumulation in macrophages at 24 hours post-infection (hpi) **(Figure 1A-B and S1A)**. To deepen this finding, we evaluated the expression of the main genes involved in lipid metabolism and LD formation in the 12 hpi. Differently from the classical LPS+IFN-γ model **(Figure S1B)**, the *E. coli* infection induced a discrete change in gene expression of lipid metabolism-associated genes at 12 hpi **(Figure 1C)**. We observed that *E. coli* infection led to a slight increase in of gene *plin2*, *dgat1* and *cd36*, and a decrease in the expression of lipolytic gene *pnpla2/atgl* **(Figure 1C)**. In comparison, stimulation with LPS+IFN-γ led to a more expressive remodeling of the expression of all genes involved in lipid metabolism **(Figure S1B)**. These findings indicate that *E. coli* infection triggers a lipid metabolic profile distinct from that induced by the classical LPS + IFN-γ.

**Figure 1:**
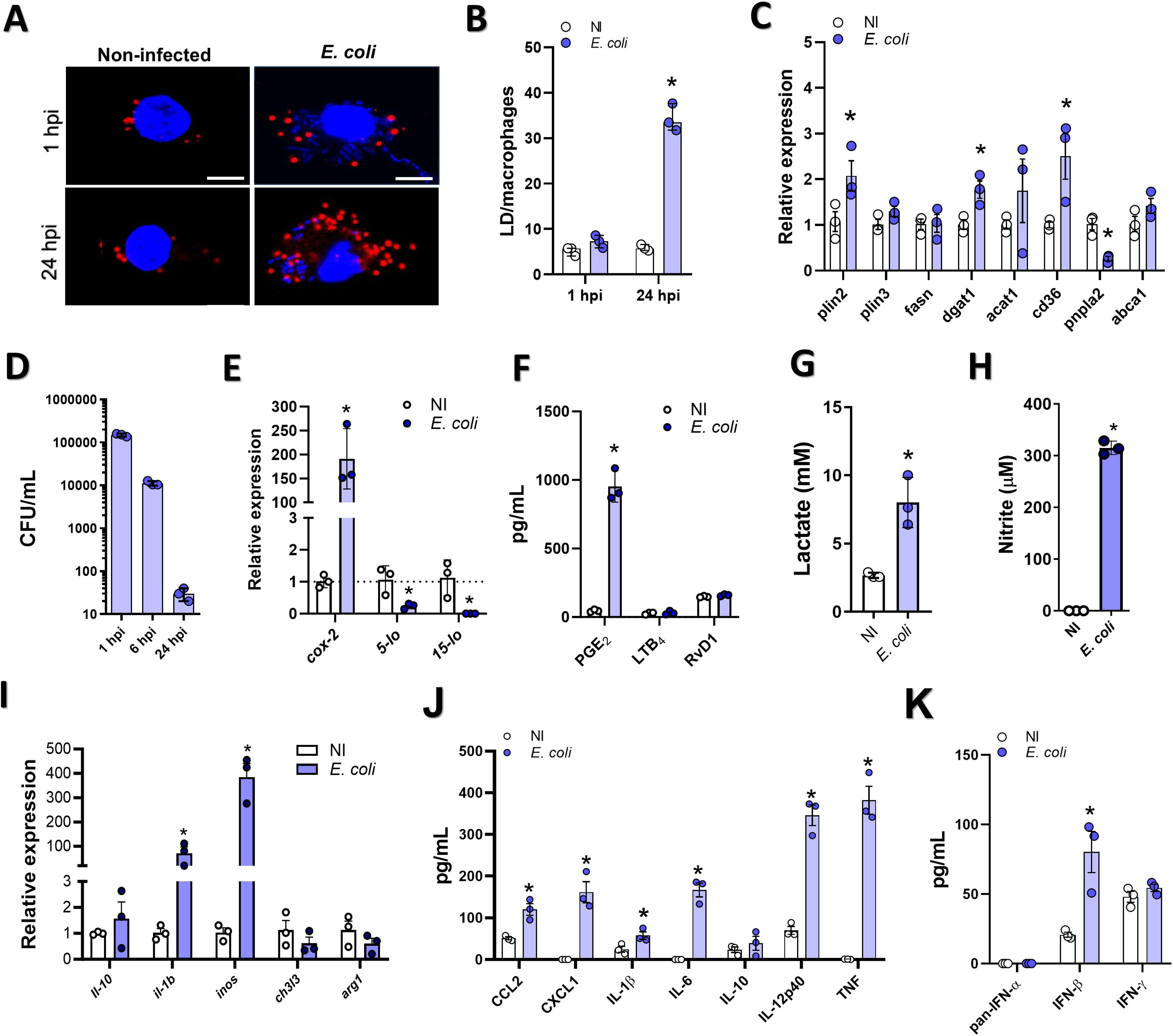
*E. coli* infection induced LD biogenesis and proinflammatory reprogramming in murine macrophages. BMDMs were infected with *E. coli* (MOI 100) for 1 hour, followed by gentamicin treatment to remove extracellular bacteria. (A) Confocal images showing Oil Red O-labeled LDs (red) in infected BMDMs. Nuclei were stained with DAPI (blue). Scale bar: 10 μm.(B) Quantification of LDs at 1 and 24 h post-infection (hpi). (C) Relative mRNA expression of lipid metabolism genes (*plin2, plin3, fasn, dgat1, acat1, cd36, pnpla2/atgl, abca1*) at 6 hpi via RT-qPCR. Data normalized to *bactin*. (D) Intracellular *E. coli* CFUs at 1 and 24 hpi. (E) mRNA expression of *ptgs-2/cox-2, 5-lo,* and *15-lo* in noninfected (NI) and infected BMDMs at 6 hpi. (F) Levels of PGE_2_, LTB_4_, and RvD1 in supernatants at 24 hpi (EIA assay). (G) Lactate levels in supernatants at 24 hpi (enzymatic assay). (H) Nitrite (NO) levels in supernatants at 24 hpi (Griess method). (I) Expression of pro-inflammatory (*il-1b, inos*) and anti-inflammatory (*il-10, ch3l3, arg1*) genes at 6 hpi via RT-qPCR. (J) Levels of cytokines (CCL2, CXCL1, IL-10, IL-1β, IL-6, IL-12p40, TNF) at 24 hpi (ELISA). (K) Levels of pan-IFN-α, IFN-β, and IFN-γ at 24 hpi (ELISA). Data represent mean ± SEM from three independent experiments. Significant difference (p < 0.05) compared to NI group.

Our next step was to evaluate whether the accumulation of neutral lipids could be associated with *E. coli* intracellular viability. Unlike pathogenic bacteria ^13,37^, *E. coli* viability decreased sharply between 1 hour and 24 hours (Figure 1D), suggesting that LD accumulation does not contribute to the intracellular survival of bacterium. To assess whether this LD accumulation would be associated with a pro-inflammatory context during *E. coli* infection. First, we evaluated the expression of the key enzymes involved in lipid mediator synthesis (*cox-2*, *5-lo*, and *15-lo*). The *E. coli* infection increases the expression of *cox-2* and decreases the expression of *5-lo* and *15-lo* **(Figure 1E).** Reinforcing this finding, we observed that the infection led to an increase in the synthesis of PGE_2_, but not LTB_4_ or RvD1 **(Figure 1F)**. Moreover, we observed that *E. coli* infection induced classical glycolytic shift **(Figure S1C and 1G)** and pro-inflammatory reprogramming in macrophages. This was evidenced by elevated nitric oxide production **(Figure 1H),** an increase in the expression of *inos/nox2* and *il-1b* **(Figure 1I)**, and higher levels of pro-inflammatory chemokines (CCL2 and CXCL1) and cytokines (IL-1b, IL-6, IL-12p40, and TNF) in the supernatant **(Figure 1J)**. There was no increase in the gene expression of M2 biomarkers (*il-10*, *arg-1*, *ch3l3*) **(Figure 1H)**, nor in the production of IL-10 **(Figure 1J)**. In host-pathogen interaction, several works have reported that LDs are essential for enhancing the synthesis of IFN in infected cells, playing a critical role in the early innate response to viral infection^38^. Our next step was to evaluate whether *E. coli* infection could induce IFN production in macrophages. Interesting, the *E. coli* infection induced the release of IFN-β but not IFN-α or IFN-γ by macrophages **(Figure 1K)**, suggesting a selective IFN response within this extracellular bacterial context.

### The modulation of lipid droplets impaired the bacterial clearance in macrophages

Our previous data in human macrophages indicated that the interaction between LD and *E. coli* was associated with a decrease in bacterial viability ^24^. In murine macrophages, LDs interact with *E. coli* (Figure 2A), but these interactions are relatively infrequent, occurring in only about 19% of cases. Bacterial interactions with LDs can promote bacterial survival or inhibit their viability depending on the bacteria and the host cell and stimulatory conditions ^5^. Using bioparticles of *E. coli*, we observed that this interaction occurred independently of bacterial viability **(Figure 2B)**. As a proof-of-principle of the protective role of LD for bacterial infection, we purified LD (**Figure 2C)** from LPS+IFN-stimulated macrophages, and we conducted a BPKA assay as reported by Bosh *et al*. (2020). We observed that LD from LPS+IFN-γ-stimulated macrophages demonstrated enhanced bactericidal activity than the control (**Figure 2D-F**).

**Figure 2:**
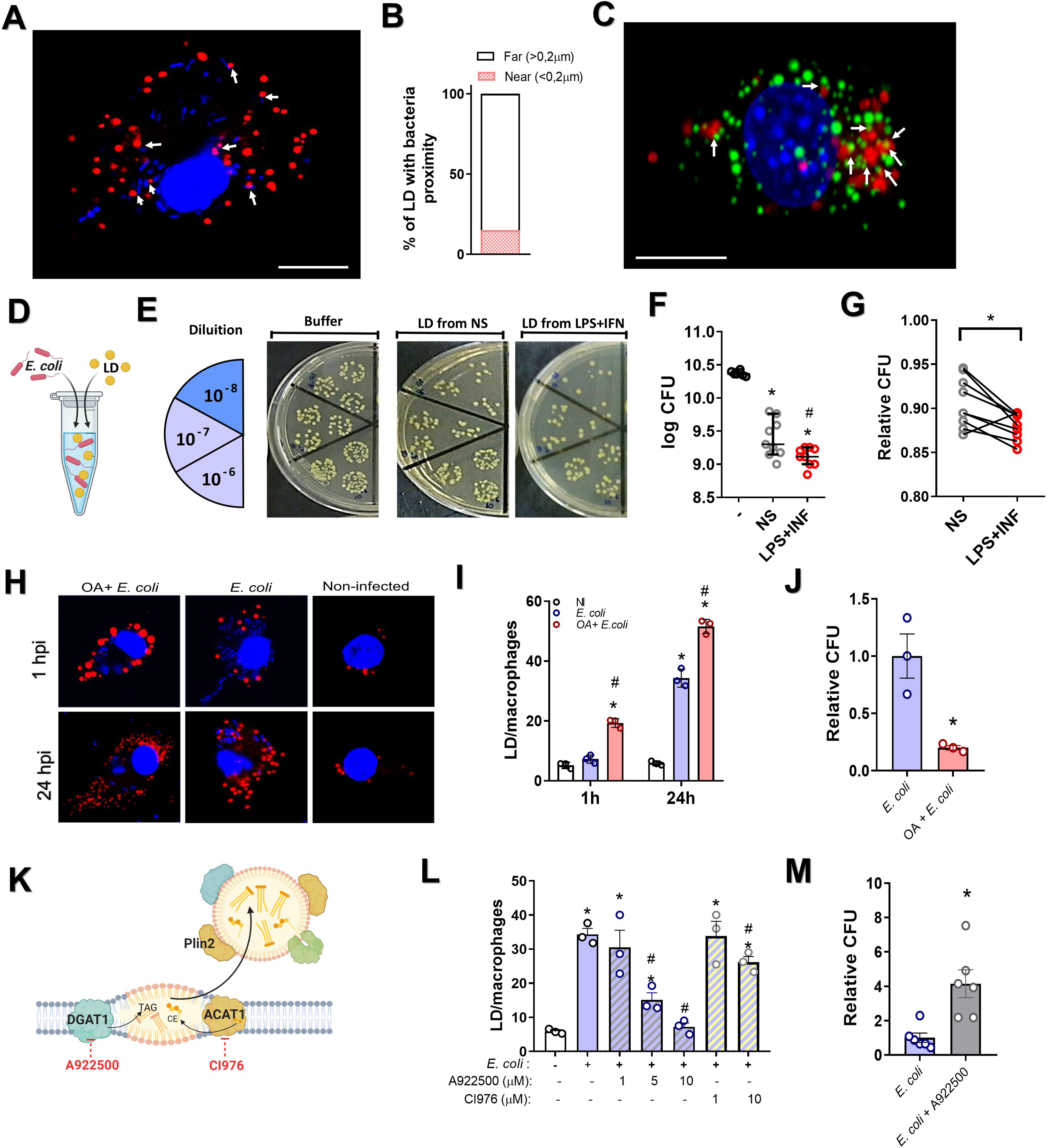
Lipid droplets contributed to antibacterial activity of macrophages. (A) Confocal images of Oil Red O-stained lipid droplets (LDs, red) in *E. coli*-infected BMDMs. Nuclei stained with DAPI (blue). Scale bar: 10 μm. (B) Quantification of the distance between LD and bacteria was performed using Fiji/ImageJ software from confocal images. (C) Confocal images of Bodipy-labeled LDs (green) in BMDMs stimulated with Texa-red *E. coli* bioparticles (red). Nuclei stained with DAPI (blue). Scale bar: 10 μm. (D) Experimental design of bacterial plate killing assays (BPKA). (E) Macrophages stimulated with LPS (500 ng/mL) and IFN-γ (10 ng/mL) for 24 h. LDs purified by sucrose gradient were tested in BPKA. (F) BPKA results using LDs from non-stimulated (NS) and LPS+IFN-γ-stimulated macrophages. (F) CFU quantification (n=9/group). (G) Relative bacterial viability compared to saline control (n=9/group). (H) Confocal images of BMDMs treated with oleic acid (OA, 40 µM) for 16 h after 1 hpi and 24 hpi. Scale bar: 10 μm. At least 100 cells across 10 fields analyzed per group. (I) Relative intracellular bacterial CFUs in macrophages with or without OA pretreatment after 24 hpi. (K-L) BMDMs infected with *E. coli* (MOI 100) with or without DGAT1 (A922500, 1.5 or 10 μM) or ACAT1 (CI976, 1 or 10 μM) inhibitors. (K) Experimental design (BioRender). (L) LD enumeration in infected and uninfected macrophages at 24 hpi (≥100 cells/10 fields/group). (M) Relative intracellular bacterial CFUs at 24 hpi. Data represent mean ± SEM from three experiments. *p<0.05 compared to non-infected controls.

Our next step was to investigate the contribution of these findings to control bacterial load into the macrophages. Our first experimental strategy was to stimulate LD biogenesis by pretreating macrophages with 40 µM oleic acid (OA) for 16 h before bacterial infection. Pretreatment with OA promoted LD biogenesis, increasing LD quantity at 1 hpi and 24 hpi **(Figure 2G-H).** We observed that OA reduced the number of viable intracellular bacteria at 24 hpi **(Figure 2I and Figure S2A).** However, the reduction in bacterial numbers at 1 hpi following OA pretreatment suggests it may also be linked to decreased bacterial uptake. Because of this limitation, our next strategy was to impair the LD accumulation by inhibiting the main pathway associated to neutral lipid accumulation **(Figure 2J)**. The treatment with DGAT1 inhibitor (iDGAT1) was able to reduce *E. coli*-induced LD accumulation **(Figure 2K)**. On the other hand, The ACAT-1 inhibitor has a minimal effect on reducing LD biogenesis. Next, we examined whether inhibiting DGAT1 could alter the levels of macrophage intracellular bacteria load. The inhibition of DGAT-1 enzyme led to an increase in the number of remaining intracellular bacteria at 24 hpi **(Figure 2L)**, and this phenomeon was not associated to alteration in bacterial uptake **(Figure S2B)**. We also evaluated whether iDGAT1 could influence *E. coli* proliferation. Our data did not indicate any change *in vitro* proliferation of this bacterium in LB medium **(Figure S2C)**. Taken together, these results suggest that changes in the lipid metabolism of macrophages affect the killing capacity of macrophages.

### Inhibition of LD accumulation reduced the inflammatory response of macrophages and impaired the IFN production and function

Next, we evaluated whether inhibition of LD accumulation by iDGAT1 treatment could influence the pro-inflammatory response induced by *E. coli* infection in macrophages. We observed that avoiding LD accumulation was associated with a significant decrease in the inflammatory lipid mediator PGE_2_ but not LTB_4_ in this model **(Figure 3A)**. Additionally, DGAT1 inhibition reduced the levels of lactate production **(Figure 3B)**. These results prompted us to analyze the activation of macrophages treated with iDGAT1. We found that inhibition did not affect the pro-inflammatory activation of macrophages, as indicated by the expression levels of F4/80 **(Figure S3A)**, CD80 **(Figure S3B)**, and GLUT1 **(Figure S3C)**. Likewise, the expression of M2-marker CD206 remained unchanged when compared to *E. coli*-untreated group **(Figure S3D)**. Interestingly, infection-induced CD36 expression was enhanced by iDGAT1 treatment **(Figure S3E)**, suggesting a potential compensatory mechanism.

**Figure 3:**
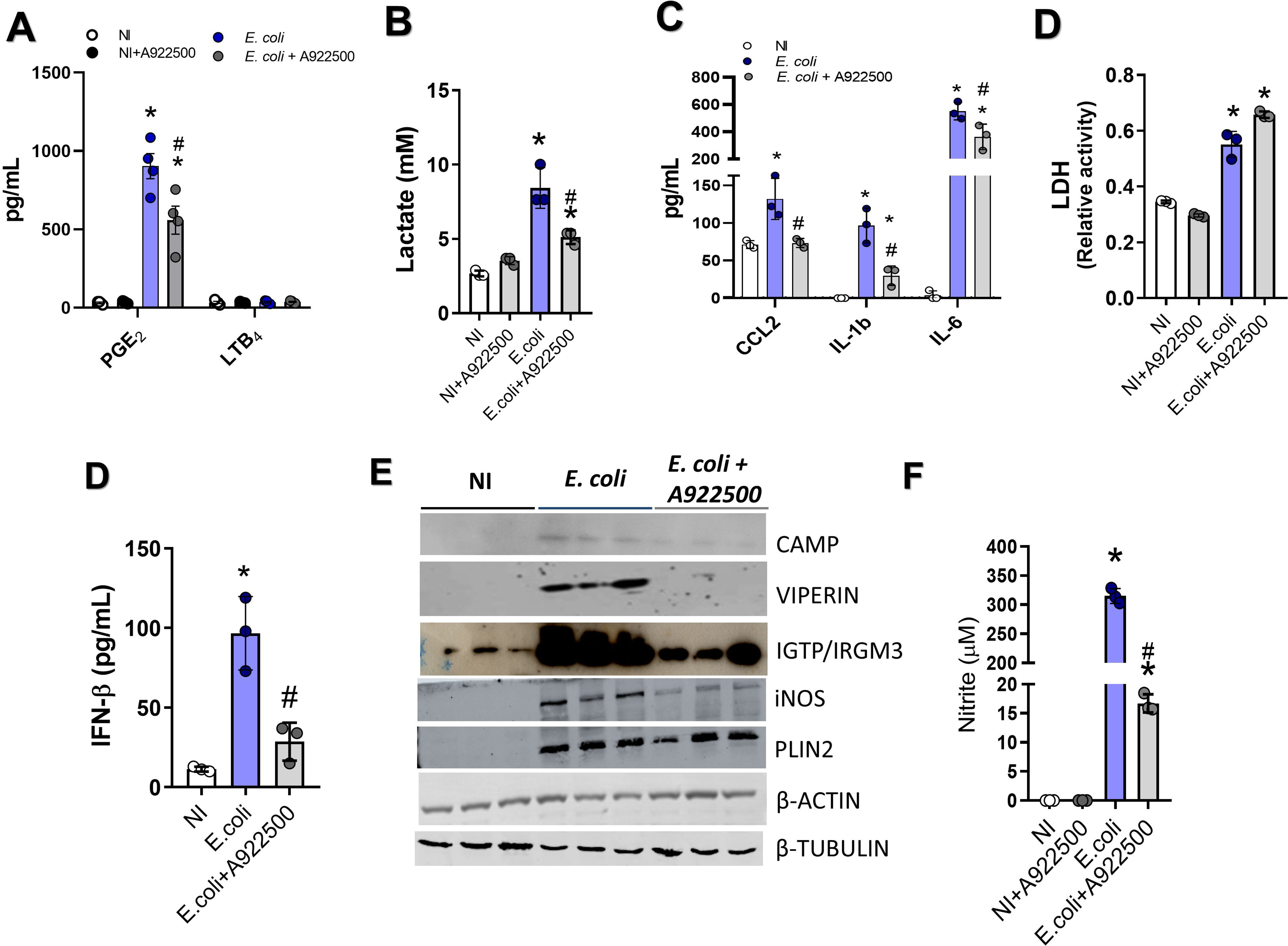
DGAT1 inhibition disrupted proinflammatory reprogramming and interferon signaling induced by *E. coli* in macrophages. BMDMs were infected with *E. coli* (MOI 100) with or without 10 μM DGAT1 inhibitor (A922500) pretreatment. (A) PGE_2_ and LTB_4_ levels in cell-free supernatants at 24 hpi were measured by EIA. (B) Lactate levels in supernatants at 24 hpi were assessed using an enzymatic assay. (C) CCL2, IL-1β, and IL-6 levels in supernatants at 24 hpi were determined by ELISA. (D) LDH release in supernatants indicated cellular viability. (E) IFN-β levels in supernatants at 24 hpi were measured by ELISA. (F) CAMP, IGTP/IRGM3, iNOS, and Plin2 expression in cell lysates were analyzed by Western blotting, using β-actin and β-tubulin as loading controls. (G) Nitrite (NO) levels in supernatants at 24 hpi were quantified by the Griess method. Data represent the mean ± SEM of three independent experiments. (*) indicates a value significantly different (p<0.05) from that of the respective noninfected group.

Next, we evaluated the impact of iDGAT1 treatment on inflammatory cytokines and chemokines production. The inhibition of LD accumulation reduced the production of CCL2, IL-1β and IL-6 induced by *E. coli* infection **(Figure 3C),** but not the lytic cell death induced by bacterial infection measured by LDH activity (**Figure 3D)**. However, no alteration was observed in *E. coli* -induced CXCL1, IL-10 IL-12p40, TNF or IFN-γ in the iDGAT1-treated group **(Figure S3F)**. These results suggest that DGAT1 pathway plays a selective role in modulating inflammatory mediators without fully deactivating the macrophage response.

Furthermore, we observed that preventing LD accumulation with iDGAT1 inhibit *E. coli*-induced IFN-β **(Figure 3D)**. Next, we investigated whether iDGAT1 treatment would impact the expression of Interferon-Stimulated Genes (ISGs), specifically Viperin, CAMP, and IRGM3/IGTP, which have previously been linked to LDs presence in host cells ^24,39,40^. The inhibition of DGAT1 in macrophages reduced the expression of CAMP, Viperin, IGTP **(Figure 3E)**, and impaired NO production, aligning with reduced INOS expression **(Figure 3E-F)**. Together, these findings suggest a significant role of DGAT1 and LD accumulation in the host signaling response critical for resistance to bacterial infection.

### The loss of DGAT1 expression decreases LD accumulation and inflammatory function in bacterial infection

To further confirm the role of DGAT1 in LD accumulation and inflammatory response during *E. coli* infection, we knocked down *dgat1* expression in murine macrophages. Initially, we tested two methods. The first approach utilized siRNA with lipofectamine, which reduced *dgat1* expression by approximately 50% **(Figure S4A)**. The second approach employed self-delivery siRNA technology, which achieved around 75% knockdown efficiency in primary macrophages **(Figure S4B-C)**. Due to its higher effectiveness, we proceeded with the self-delivery siRNA method. Knockdown of *dgat1* reduced *E. coli*-induced LD accumulation **(Figure 4A-B)** and increased bacterial persistence but did not affect phagocytosis **(Figure 4C and S4D)**. In alignment with our pharmacological inhibition findings, reducing DGAT1 expression decreased the production of PGE_2_ **(Figure 4D),** and the levels of IL-6, CCL2, and IFN-β **(Figure 4E-G)**. DGAT1 downregulation also lowered NO production **(Figure 4H)**. We found that DGAT1 loss corresponded with reduced *inos* and *cox2* expression **(Figure 4I-J)**. Unlike pharmacological DGAT1 inhibition, DGAT1 knockdown did not impact lactate production **(Figure 4K)** or LDH release **(Figure 4L)**. Collectively, these results confirm the central role of DGAT1 in LD accumulation and in regulating bacterial activity in murine macrophages.

**Figure 4:**
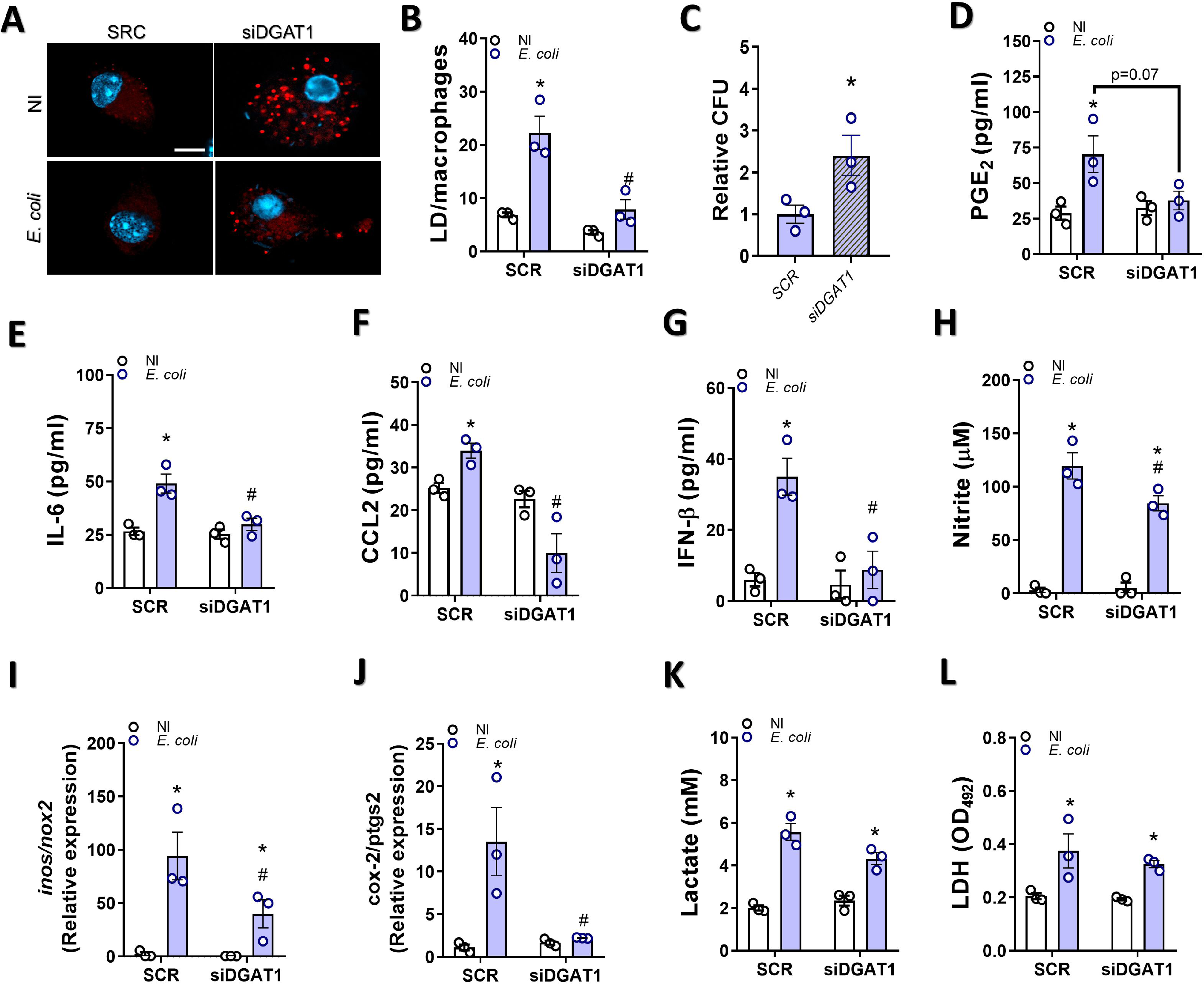
RNAi-mediated DGAT1 knockdown impaired antibacterial activity, proinflammatory reprogramming, and interferon signaling in E. coli-infected macrophages. BMDMs were transfected with 5 µM self-deliverable AUMsilence oligos for DGAT1 or scramble RNA (SCR). After 48 h, BMDMs were infected with *E. coli* (MOI 100). (A) Oil Red O-stained LDs (red) in infected BMDMs at 24 hpi, with nuclei stained by DAPI (blue). Scale bar: 10 μm. (B) LD quantification in infected BMDMs (≥100 cells per group across 10 fields/experiment). (C) Relative intracellular bacterial CFU at 24 hpi. (D) PGE_2_ levels in supernatants at 24 hpi by EIA. Levels of (E) IL-6 and (G) IFN-β in supernatants at 24 hpi by ELISA. (H) Nitrite (NO) levels in supernatants at 24 hpi by the Griess method. (I, J) Relative mRNA expression of *inos/nox2* and *ptgs-2/cox-2* genes in noninfected (NI) and infected BMDMs at 6 hpi, normalized to *β-actin* (mean 2−ΔΔCt ± SEM; n = 3). (K) Lactate levels in supernatants at 24 hpi by enzymatic assay. (L) Cellular viability assessed by LDH release in supernatants. Data represent the mean ± SEM of three independent experiments. (*) indicates a significant difference (p < 0.05) compared to the respective noninfected group or SCR E. coli-infected group.

### DGAT1 inhibition prevents sepsis-induced LD accumulation and downregulates the protective inflammatory response

Our next step was to investigate whether DGAT1 inhibition blocked sepsis-induced LD accumulation in peritoneal leukocytes during the early stages of sepsis. In 6h and 24h post-sepsis, the DGAT1 inhibition reduced sepsis-induced LD accumulation in peritoneal leukocytes **(Figure 5A-C).** During sepsis, LDs have been shown to serve as a platform for the synthesis of lipid inflammatory mediators in leukocytes ^36,41^. We observed that inhibition of LD accumulation significantly reduced the production of lipid inflammatory mediators, including PGE_2_, LTB_4_ and RvD1 **(Figure 5D)**. These data reinforce the central of LD in the production of lipid inflammatory mediators in sepsis.

**Figure 5:**
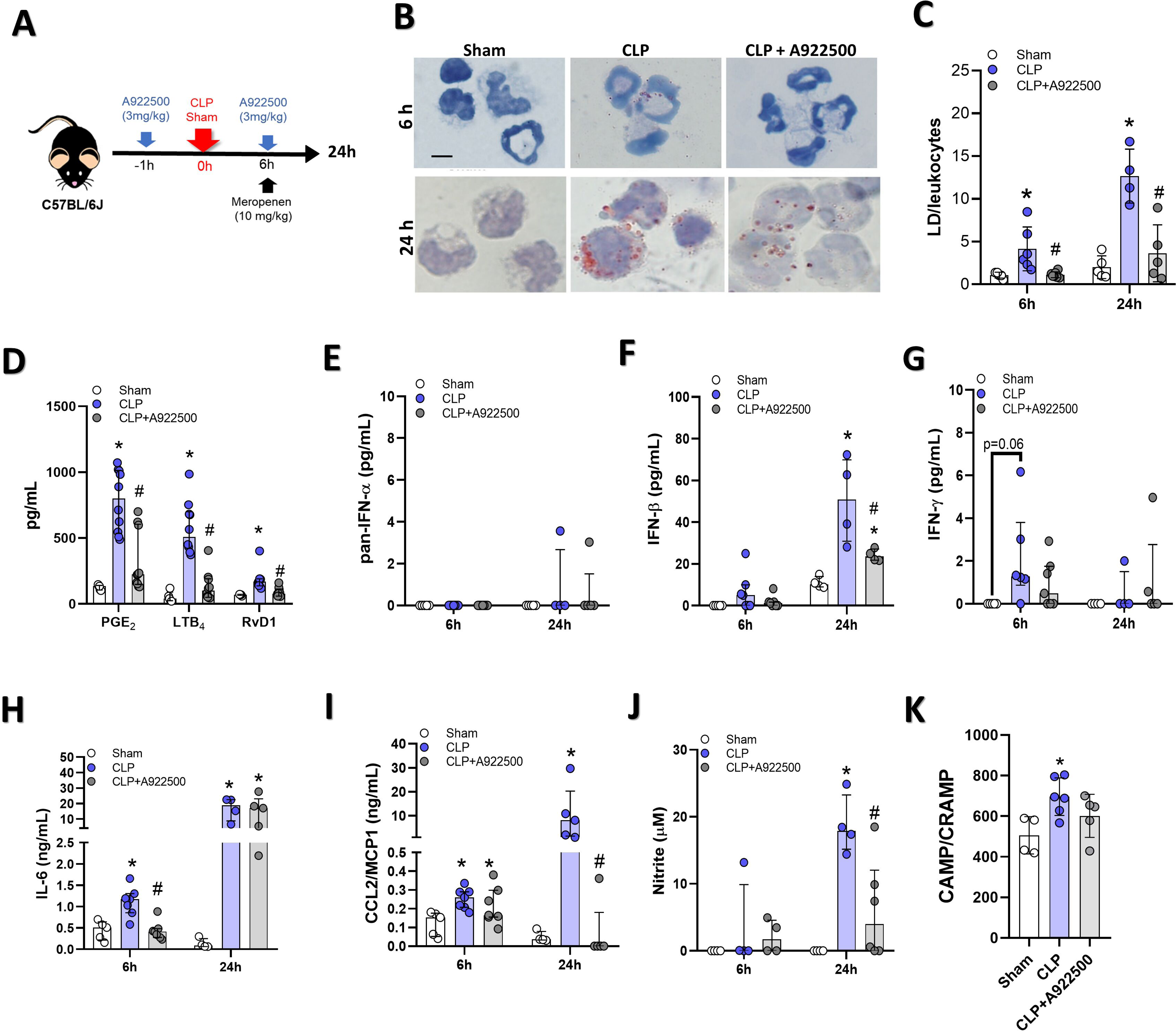
DGAT1 inhibition reduces proinflammatory responses in the peritoneum of septic mice. Sham or septic (CLP) mice were orally treated with A922500 (DGAT1 inhibitor, 3 mg/kg) or vehicle. At 6 h or 24 h post-surgery, peritoneal lavage was collected. (A) In vivo experimental design for DGAT1 inhibition. (B) Oil Red O-stained lipid droplets (red) in peritoneal leukocytes, counterstained with Mayer’s hematoxylin. Scale bar: 20 μm. (C) Lipid droplet enumeration in peritoneal leukocytes (≥100 cells/group across 10 fields/experiment). (D) PGE_2_, LTB_4_, and RvD1 levels in peritoneal lavage at 24 h post-surgery by EIA. Levels of (E) pan-IFN-α, (F) IFN-β, (G) IFN-γ, (H) IL-6, (I) CCL2, (J) nitrite, and (K) cathelicidin (CAMP) in peritoneal lavage at 6 h and 24 h post-surgery by ELISA. Data (4–6 mice/group) are presented as means ± SEM, analyzed by one-way ANOVA with Tukey’s post hoc test. *p < 0.05 compared to sham; #p < 0.05 compared to untreated CLP. CLP: cecal ligation and puncture.

We further investigated the effect of lipid droplet (LDs) inhibition on other inflammatory mediators in vivo (Figure 5E-K). Sepsis-induced LD shows an upregulation of several ISGs, including Viperin and CAMP, indicating elevated systemic interferon levels **(Figure S5A)**. We observed that the polymicrobial sepsis did not affect the production of IFN-α **(Figure 5E)**, but increased the production of IFN-β in 24h **(Figure 5F)**, accompanied by a modest and transient elevation in IFN-γ levels at 6h post-sepsis **(Figure 5G)**. The downregulation of LD accumulation induced by the IDGAT1 inhibition during sepsis was followed by a significant decrease in IFN-β levels 24h post-sepsis **(Figure 5F)**. We also observed that prevention of LD accumulation also impaired sepsis-induced IL-6 and CCL2 in 6h and 24h post-sepsis, respectively **(Figure 5H-I)**. No significant differences were observed between untreated and treated sepsis groups in IL-1β, IL-12p40, IL-10, CXCL1 and TNF production **(Figure S5B-F)**. Due to reduced CCL2 levels, we also investigated whether inhibiting DGAT1 could impair the activation of peritoneal macrophages. However, our results showed that the treatment did not reduce the pro-inflammatory activation macrophages marker (F4/80) **(Figure S5G)**. There was an increase in CD80 and GLUT1 expression in cells derived from the CLP-iDGAT1 treated group, but no changes in CD206 expression were observed between the groups **(Figure S5H-J).** Our next step was to evaluate whether DGAT1 inhibition could impair the antibacterial mechanisms during infection. We found that preventing LD accumulation also reduced sepsis-induced increase of NO **(Figure 5J)** and impaired secretion of antibacterial peptide cathelicidin (CAMP) **(Figure 5K)**. Altogether, these data support the role of DGAT and lipid accumulation in leukocyte inflammatory response during sepsis.

### The inhibition of DGAT1 led to the loss of control of the bacterial load in the sepsis

The impairment of resistance mechanism compromises the host’s ability to control infections ^27,42^. In this context, we investigated the bacterial load both in blood and in the peritoneal site in the sepsis. Bacterial load was significantly higher in the peritoneum of septic mice treated with iDGAT1 compared to untreated mice at both 6h and 24h post-sepsis **(Figure 6A)**. Moreover, the bacterial load was also significantly elevated in the serum of iDGAT1-treated mice at both times analyzed **(Figure 6B)**, indicating increased bacterial dissemination and a loss of control over the infection. To determine whether the loss of bacterial control was associated with impaired leukocyte recruitment, we also evaluated the cell number in the peritoneal cavity. However, no significant differences were observed in the total or differential leukocyte counts between untreated and treated sepsis groups at either 6 hours **(Figure 6C)** or 24 hours post-sepsis **(Figure 5D)**. Our next step was to evaluate whether LDs also had a protective role *in vivo*, in the polymicrobial sepsis model.

**Figure 6:**
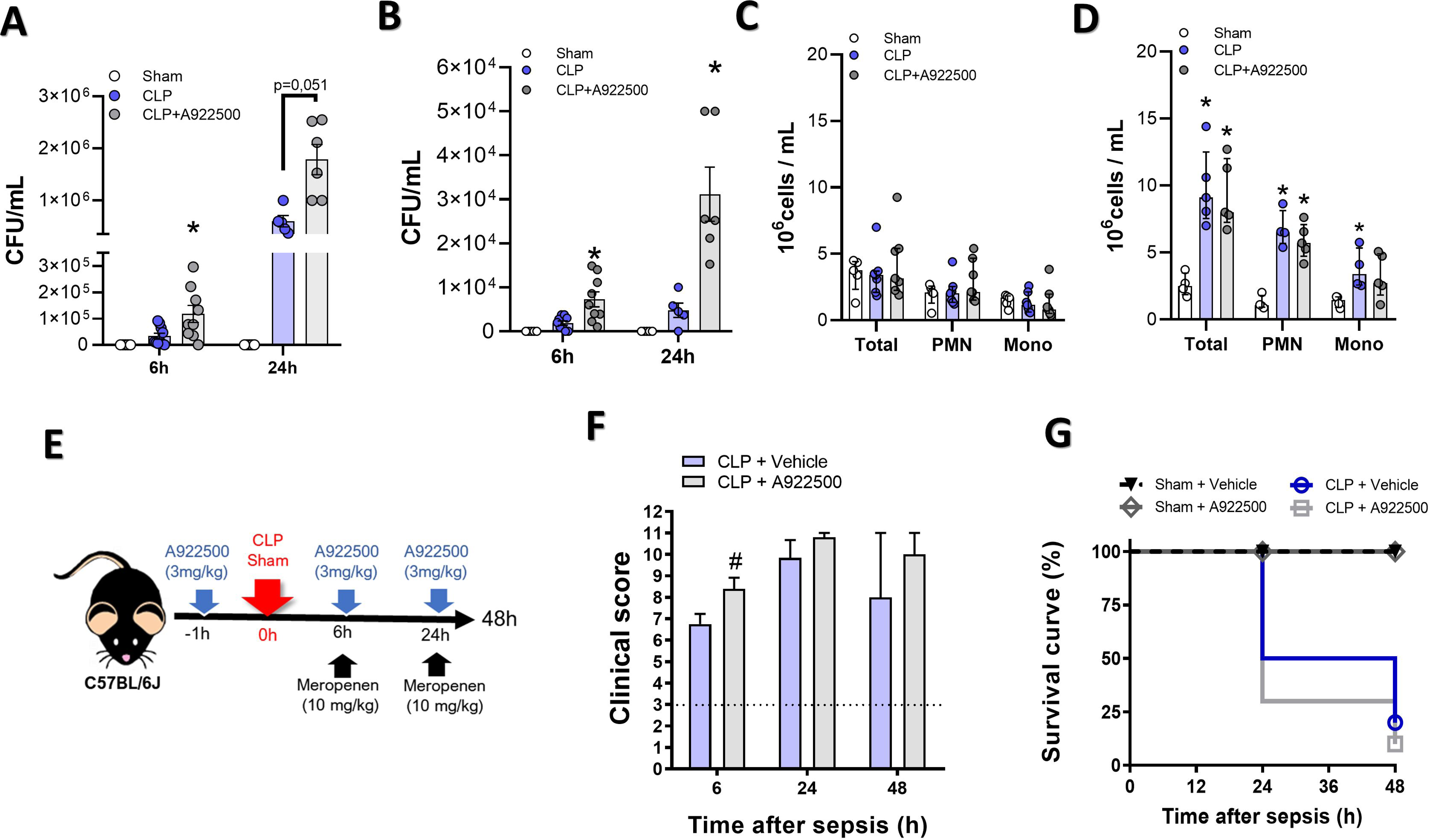
DGAT1 inhibition increases bacterial load and sepsis severity. Sham or septic (CLP) mice were orally treated with A922500 (DGAT1 inhibitor, 3 mg/kg) or vehicle. (A-B) Bacterial CFU quantification in peritoneal lavage (A) and serum (B) at 6 h and 24 h post-surgery. (C-D) Total and differential leukocyte counts in peritoneal lavage at 6 h (C) and 24 h (D), including total leukocytes, mononuclear cells (Mono), and polymorphonuclear leukocytes (PMN). (E-G) Experimental design for survival and clinical score analysis. Mice were monitored for 48 h to assess (F) clinical sepsis severity scores and (G) survival. Data (4–6 mice/group for A-D; 10 mice/group for E-G) are presented as means ± SEM, analyzed by one-way ANOVA with Tukey’s post hoc test. *p < 0.05 compared to sham; #p < 0.05 compared to untreated CLP. CLP: cecal ligation and puncture.

The pretreatment with iDGAT1 exacerbated clinical conditions at 6 hours post-sepsis, which remains extremely high at all times analyzed **(Figure 6F).** Moreover, DGAT-1 inhibition temporally anticipates the mortality associated with severe sepsis **(Figure 6G)**. Together, our results collectively suggest that LD accumulation plays a crucial role in enhancing the antibacterial properties and inflammatory response of macrophages during sepsis.

## 3. DISCUSSION

LDs have been reported in infectious disease across all pathogen classes, from viruses ^43,44^ and protozoa ^45,46^, to bacteria ^10,15^ and fungi ^47^. Host LDs are often exploited by specialized pathogens to evade the immune system and/or serve as an energy source for their survival and/or replication ^18–20^. However, recent findings highlight a drastic transformation in the participation of LDs within infectious and inflammatory contexts. Emerging research suggests that LDs support host defense ^18–20,48^, acting as regulators of immunometabolism ^24^ and essential platforms for producing protective eicosanoids, cytokines and antibacterial peptides ^41,48^. Additionally, neutral lipid synthesis and LD accumulation significantly enhance the pro-inflammatory profile of macrophages ^49,50^

Similar to the classic LPS model, *E. coli* infection induces LD accumulation in macrophages, which is associated with glycolytic reprogramming ^50–52^. This *E. coli*-induced LD accumulation is a multifaceted process primarily linked to increased lipid uptake (via CD36), reduced lipolysis (via ATGL), and elevated expression of LD structural proteins. These findings align with those of Feingold *et al.* (2012), who showed that TLR activation promotes neutral lipid accumulation in macrophages through multiple pathways ^51^. However, *E. coli*-induced LD accumulation involves minimal changes in the expression of lipid synthesis enzymes. In fact, *de novo* lipid synthesis is not required for the LD formation of classical activated macrophages ^53^.

LD biogenesis is strongly associated with PAMP recognition by TLRs in leukocytes ^8,11^, with different TLRs being engaged depending on the specific pathogen ^8,10^. In the pathogen-host context, the host response may be closely linked to the varied functions of LDs. Castoldi *et al.* (2020) reported that triglyceride synthesis and LD accumulation significantly enhance the pro-inflammatory profile of LPS-estimated macrophages, primarily by serving as a platform for PGE_2_ synthesis ^50^. These results agree with what we observed in *E. coli* infection and in sepsis, where blocking LD accumulation resulted in decreased PGE_2_ production and attenuation of the inflammatory response, without deactivating macrophages, but was associated with an increase in bacterial load. Our data highlights further the complexity of PGE_2_’s role in inflammation and infection both in vitro and in vivo. Notably, PGE_2_ can be either pro-inflammatory or anti-inflammatory depending on the pathogen, the infection site and stage, and the concentration of this mediator at the infection site ^54–58^. During *S. typhimurium* infection, PGE_2_ has also been identified as an inducer of glycolytic reprogramming and the pro-inflammatory response in macrophages ^57^. On the other hand, lactate also stimulates PGE_2_ synthesis ^59^, which can inhibit β-oxidation and lead to LD accumulation ^60^. Additionally, PGE_2_ regulates iNOS in macrophages, acting as either an activator or repressor, depending on its concentration ^61^.

In this pro-host context, the protective role of LD in infection has been associated with IFN response ^23,38,62^. However, the role of type I IFNs in bacterial infections remains less understood compared to their role in viral infections^63,64^. Our results demonstrate that preventing LD accumulation through DGAT1 inhibition impairs IFN-β release, reducing the expression of key antibacterial proteins, including iNOS and cathelicidin, which leads to an increase in bacterial load and sepsis-induced mortality. It is important to emphasize that the increase in bacterial load observed in vivo is not attributed to sepsis-induced impairment of leukocyte migration to the infection site. The results are consistent with Mancuso et al. (2006), who reported that IFN-α/β signaling is essential for host defense against various bacteria, including *E. coli* ^65^. In the absence of IFN-α/β signaling, a marked reduction in macrophage production of NO, and TNF-α was observed after stimulation with live bacteria or with purified LPS ^65^. Moreover, IFN-β also mediate time-dependent increases in the mRNA levels of microsomal PGE synthase-1 and COX-2, and induced PGE_2_ production ^66^. In the polymicrobial sepsis model, mice deficient in IFN-α/β receptor (IFNAR) display persistently elevated peritoneal bacterial counts compared with wild-type mice ^67^. Furthermore, Bosch et al. (2020) reported that LPS triggered remodeling of the LD proteome, leading to an increase in several innate immune proteins, many of which belonged to the ISG family, including Viperin, IGTP and CAMP. Interestingly, we observed that inhibiting LD accumulation not only reduced IFN-β release but also suppressed ISG expression. Our findings align with recent evidence demonstrating that type I IFN production and signaling is strongly dependent on cellular metabolism ^68,69^.

On the other hand, the role of LD as an enhancer of inflammatory response also can be a two-edged sword. In sepsis, tissue damage results from a maladaptive inflammatory and metabolic response mounted to resist infection; this is associated with inadequate mechanisms of tissue tolerance ^70^. Recently, our group demonstrated that inhibiting hepatic LD accumulation by targeting the enzyme DGAT1 reduces levels of inflammatory mediators and lipid peroxidation while improving liver function in sepsis^29^. The exacerbation of mechanisms of resistance to infection has been associated with tissue damage, which ultimately can lead to multiple organ failure ^27,28^. However, these same mechanisms involved in organ failure are essential to control bacterial loading, such as LTB_4_, NO, and CCL2 ^71,72^. The same phenomenon has been reported to IFN-β, its effects during bacterial infections can be either protective or detrimental, depending on the specific bacterium and host status ^73,74^.

In summary, our data suggests that LD biogenesis in *E. coli*-infected macrophages plays a critical role in controlling bacterial load and enhancing the innate immune response. In sepsis, preventing LD accumulation through DGAT1 inhibition disrupts the production of inflammatory mediators and antibacterial factors, leading to increased bacterial load and higher sepsis-associated mortality. These findings highlight LD accumulation as a component of cellular metabolic reprogramming and a molecular switch in innate immunity. Given the growing resistance to current antibiotics, this study provides insights into the molecular mechanisms underlying antimicrobial defense, which could inform the development of novel anti-bacterial strategies.

## Acknowledgments

The authors are grateful to the Microscopy Facility of the Brazilian National Cancer Institute for the acquisition of confocal images; and to FIOCRUZ Luminex Platform (*RPT03C* Rede de Plataformas PDTIS, FIOCRUZ/RJ) and the assistance of Edson F. de Assis for the use of Luminex facilities.

## Author Contributions

FP-D and PB conceptualized the study. FP-D was responsible for design and conducted the experiments, literature review, prepared the figures and tables. The manuscript was written by FP-D and PB, and edited by all authors. JCS, RVS, HE-S, TSS, TC-F, FFM, LP, MMC, DMO, VCS and LS-M contributed to the *in vitro* data collection and analysis. EKS and MAJ contributed to microscopy data collection and analysis. GI and MFSC contributed to cytometry data collection and analysis. FP-D LT, TC-F, TI, and PAR performed *in vivo* treatment and infection. LT, LS-M, MFSC, PAR, and PB contributed to study design and critically revised the article. All authors approved the submitted version.

